# Elucidating the mechanism of *Trypanosoma cruzi* acquisition by triatomine insects: evidence from a large field survey of *Triatoma infestans*

**DOI:** 10.1101/2020.03.23.002519

**Authors:** Aaron W. Tustin, Ricardo Castillo-Neyra, Laura D. Tamayo, Katty Borini-Mayorí, Renzo Salazar, Michael Z. Levy

## Abstract

Blood-sucking triatomine bugs transmit the protozoan parasite *Trypanosoma cruzi*, the etiologic agent of Chagas disease. We measured the prevalence of *T. cruzi* infection in 58,519 *Triatoma infestans* captured in residences in and near Arequipa, Peru. Among bugs from infected colonies, *T. cruzi* prevalence increased with stage from 12% in second instars to 36% in adults. Regression models demonstrated a linear relationship between infection prevalence and developmental stage. Prevalence increased by 5.4 percentage points with each additional stage. We postulate that the probability of acquiring the parasite may be related to the number of feeding events. Transmission of the parasite does not appear to be correlated with the amount of blood ingested during feeding. Similarly, other hypothesized transmission routes such as coprophagy fail to explain the observed pattern of prevalence. Our results could have implications for the feasibility of late-acting control strategies that preferentially kill older insects.

## 1. Introduction

*Trypanosoma cruzi*, the protozoan parasite that causes Chagas disease, is transmitted via the feces of blood-sucking triatomine insects (Hemiptera: Reduviidae) such as *Triatoma infestans*, the major vector of Chagas disease in much of South America [1]. Triatomines pass through five nymphal instar stages before becoming adults. All triatomine stages are at risk for acquiring *T. cruzi* because all of them ingest blood meals. Insects may also become infected when they engage in behaviors such as coprophagy [2]. Once acquired, infection with *T. cruzi* is persistent [3]. Therefore, in well-stablished colonies with stable transmission patterns, *T. cruzi* prevalence is expected to increase monotonically with the insects’ developmental stage. The actual shape (e.g., linear, exponential, logarithmic) of the rising prevalence curve may give clues as to the method of transmission. With respect to Chagas disease control, it may be important to ascertain the shape of this relationship between *T. cruzi* prevalence and stage.

In this paper we report the stage-prevalence of *T. cruzi* in a large sample of *T. infestans* captured during vector control and surveillance activities in Arequipa, Peru. These data allow us to test three hypotheses of *T. cruzi* acquisition by triatomines. Our first hypothesis, hereafter called the “Blood Hypothesis,” is that insects become infected with *T. cruzi* in proportion to the amount of blood they ingest. Using data from Rabinovich [4], we found that the cumulative blood ingested by *T. infestans* increases exponentially with stage (see Appendix A). If the Blood Hypothesis is correct, there should be an exponential rise in *T. cruzi* prevalence with stage (Figure 1A), with a growth constant comparable to the estimate from the feeding data. A second possibility is that an insect’s risk of infection is proportional to the number of opportunities to acquire the parasite (“Bites Hypothesis”), rather than the amount of blood ingested. Under this hypothesis, *T. cruzi* prevalence among instars would likely rise in a linear fashion with stage (Figure 1B), as all nymphs probably take roughly equal numbers of bites. The third hypothesis is that triatomines frequently acquire *T. cruzi* via coprophagy (“Coprophagy Hypothesis”).

**Figure 1.**
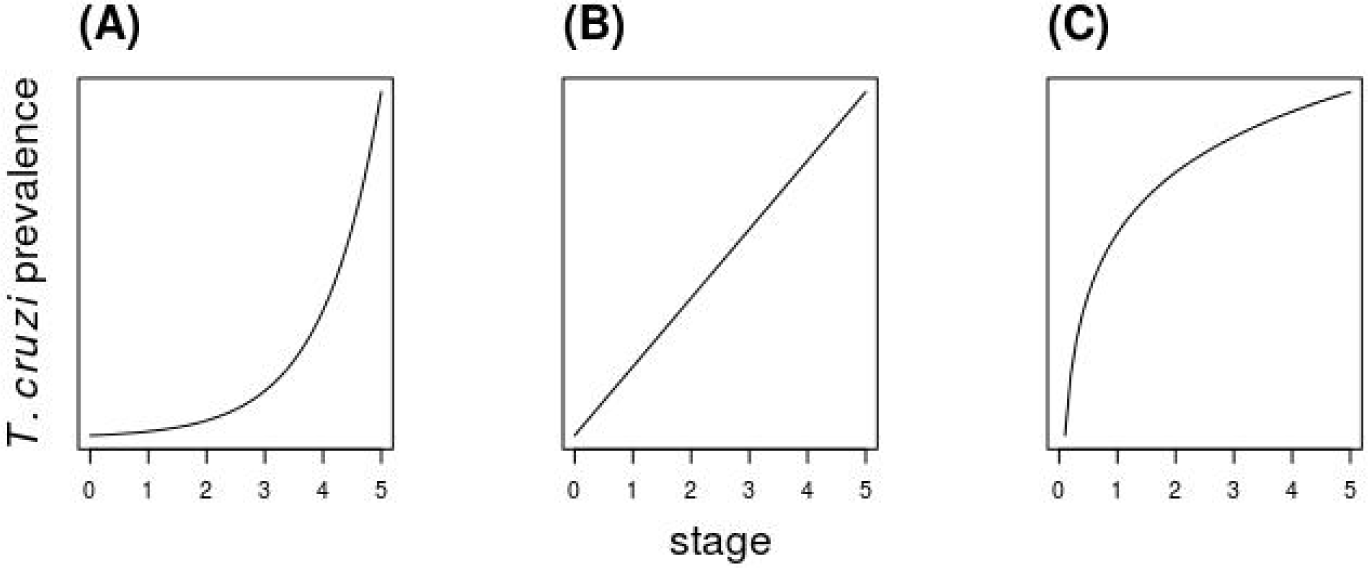
Hypothetical relationships between triatomine stage and prevalence of infection with *Trypanosoma cruzi*. (**A**) Acquisition of the parasite is proportional to volume of blood ingested (“Blood Hypothesis”); (**B**) Parasite acquisition is proportional to the number of exposure opportunities (“Bites Hypothesis”); (**C**) Early instar nymphs frequently acquire the parasite via coprophagy, while older instars acquire the parasite at a lower rate via ingestion of blood from mammalian hosts (“Coprophagy Hypothesis”).

Triatomines ingest small amounts of feces of older insects in order to acquire bacterial symbionts necessary for digestion of blood meals [5]. Presumably, it is the early instars that ingest feces, as nymphs of some species fail to mature in the absence of symbionts [6]. An isolated experiment has suggested that *T. infestans* may also acquire *T. cruzi* by this route [2]. If parasite transmission via coprophagy is common under field conditions, then *T. cruzi* prevalence may increase relatively quickly among first and second instars, followed by a less rapid increase in older nymphs that acquire the parasite only via blood feeding (Figure 1C).

## 2. Materials and Methods

Between 2008 and 2014, workers from the Arequipa Ministry of Health and the Universidad Peruana Cayetano Heredia/University of Pennsylvania (UPCH/Penn) Zoonotic Disease Research Laboratory collected triatomines from urban and peri-urban households in and around the city of Arequipa, Peru. The collection methods have been described previously [7,8]. All insects were categorized by sex (for adults), stage, and site. A site was defined as all rooms and peri-domestic areas, such as animal enclosures, associated with a single dwelling. First instar insects did not undergo further analysis. The remaining triatomines were inspected for *T. cruzi* at the UPCH/Penn Zoonotic Disease Research Center in Arequipa, using standard techniques [9]. Briefly, we extracted feces by applying pressure to each insect’s abdomen with forceps. Feces were diluted in 10 μL normal saline and examined microscopically at 400x magnification for the presence of parasites.

We built regression models to explore the relationship between *T. cruzi* prevalence and developmental stage. Due to uncertainty surrounding the total number and size of blood meals taken by adult insects, we excluded adults from our analysis. To avoid biasing our results by including a large number of insects with no potential exposure, we also excluded vectors captured at sites with no infected triatomines. We assessed the Bites Hypothesis by performing simple linear regression of infection status on stage. We used a likelihood ratio test to compare an unconstrained linear model to a second linear model in which the regression line was forced to pass through the origin (i.e., intercept = 0) to account for the fact that unfed insects are uninfected. We tested the Blood Hypothesis by regressing *T. cruzi* infection status on an exponential function of stage. We fixed the growth constant of the exponential function at k=1.33 (see Appendix A for derivation) and fitted the scale factor as an unknown model parameter. In a sensitivity analysis, we varied the value of the exponential model’s growth constant from k = 0.67 to k = 2.7 (i.e., from half to twice our point estimate). This sensitivity analysis was motivated by the fact that triatomines in our sample may have taken fewer or more blood meals than we presumed, and actual blood meal sizes may have differed from the laboratory-derived values that informed our primary analysis [10]. Finally, because the mathematical relationship between feces ingestion and infection transmission is unknown, we did not create a specific model to test the Coprophagy Hypothesis. Instead, we made a qualitative assessment of the importance of feces-mediated transmission by examining whether the other two models underestimated the infection prevalence in early-stage insects. We used the Akaike information criterion (AIC) [11] to guide model selection. Analyses were performed in R version 3.0.2 (R Foundation for Statistical Computing, Vienna, Austria).

## 3. Results

We captured and analyzed 58,519 triatomines from 4,138 sites. There were 188 sites (4.5%), harboring 15,252 insects, with at least one *T. cruzi*-infected triatomine. The stage distribution and *T. cruzi* prevalence of captured insects varied considerably between sites (Figure 2). Second instars were underrepresented at most sites, possibly because it is difficult to find and capture these small nymphs. The population structure of third instars through adults was flat in some colonies (e.g., Figure 2A and 2C), while other colonies were skewed toward younger (Figure 2F) or older (Figure 2D) insects. These differences may be due to the age of the colonies, or they may represent heterogeneity, across sites, in stage-dependent survival. *T. cruzi* prevalence was high in some colonies (e.g., Figure 2E and 2I), while other colonies had only one or a few infected insects (e.g., Figure 2 B, C, and D). We speculate that the colonies with very few infected insects may represent sites where *T. cruzi* was recently introduced, perhaps via a newly infected host or migration of an infected triatomine from a nearby household. Averaged across all 188 sites, the prevalence of *T. cruzi* infection rose monotonically from 12% in second instars to 36% in adults (Table 1 and Figure 3).

**Table 1.** *Trypanosoma cruzi* infection status by developmental stage, for 15,252 triatomines captured in 188 infected colonies in Arequipa, Peru.

| Stage | Total number of insects | Number infected | *T. cruzi* prevalence (%) |
| --- | --- | --- | --- |
| Second instar | 1,037 | 125 | 12.1 |
| Third instar | 3,352 | 512 | 15.3 |
| Fourth instar | 3,310 | 687 | 20.8 |
| Fifth instar | 3,838 | 1,060 | 27.6 |
| Adult | 3,715 | 1,326 | 35.7 |
| Total | 15,252 | 3,710 | 24.3 |

**Figure 2.**
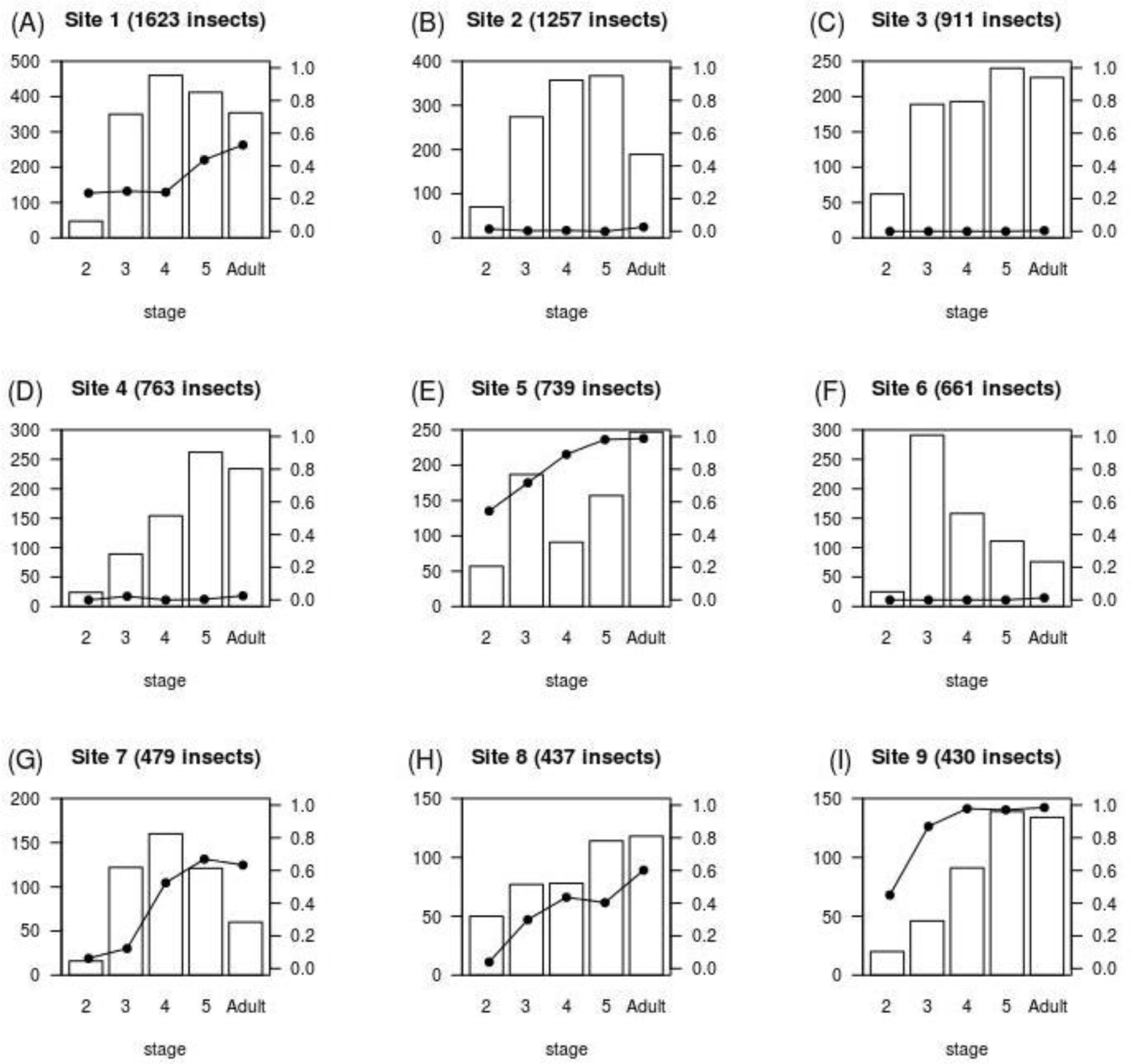
Distribution of developmental stage and *Trypanosoma cruzi* infection status in *Triatoma infestans* from nine households in Arequipa, Peru. Shown are the nine infected sites with the largest number of captured insects. White black bars and left vertical axis labels: total number of insects. Black circles and right vertical axis labels: fraction of insects with *T. cruzi*.

**Figure 3.**
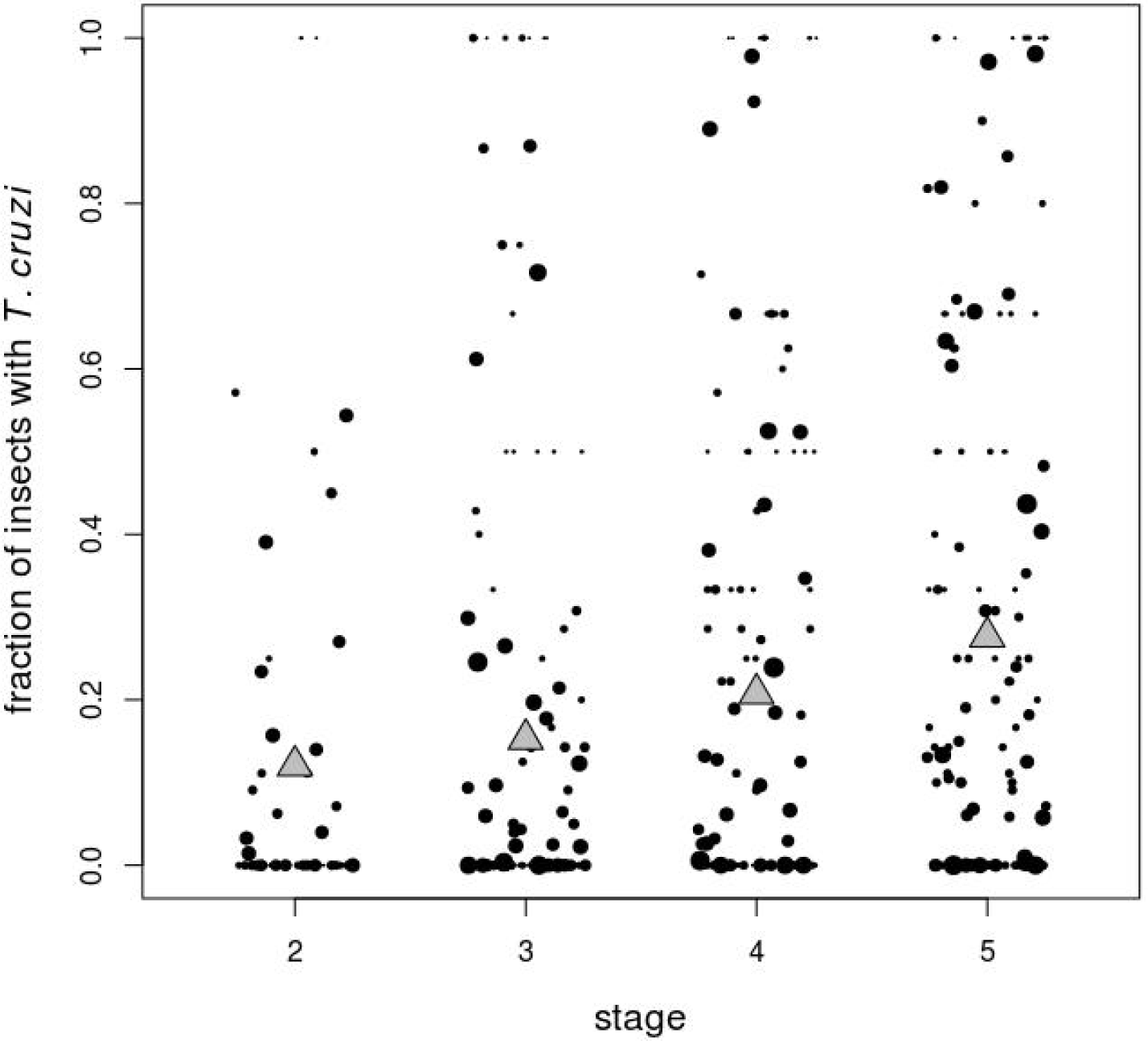
Fraction of nymph *Triatoma infestans* infected with *Trypanosoma cruzi* at 188 sites with at least one infected insect. Black circles represent individual sites; sizes of circles are proportional to the logarithm of the number of insects captured. Gray triangles show the mean fraction of insects with *T. cruzi* across all infected sites.

We found a strong linear relationship between *T. cruzi* prevalence and instar stage (Figures 3 and 4). There was no evidence of the exponential increase in prevalence that would occur if parasite transmission depended on cumulative blood ingested, nor did the prevalence curve exhibit the nonlinear behavior (i.e., excess prevalence in early nymphs) that would occur if coprophagy were a primary driver of infection. Regression modeling confirmed the linear association between prevalence and stage. The linear model fit the data well and had the lowest AIC (Table 2 and Figure 4). The likelihood ratio test did not favor the inclusion of a non-zero intercept in the linear model (p = 0.47 for comparison of null and extended models). The exponential model was a very poor fit and had a significantly higher AIC than that of the linear model. The exponential model remained a much worse fit than the linear model when we varied the growth factor parameter (range of AIC of exponential models: 11925 to 12890).

**Table 2.** Results of regression modeling. The linear models are simple linear regressions of *T. cruzi* infection status on stage. The exponential model is a nonlinear regression that assumes infection status is proportional to an exponential function of stage, with growth constant (k=1.33) informed by prior observations of *T. infestans* blood meal sizes.

| Model | Coefficient (95% confidence interval) |  |  | AIC | LRT |
| --- | --- | --- | --- | --- | --- |
|  | Stage | Intercept | Scale factor |  |  |
| Linear model | 0.056 (0.049, 0.064) | -0.011 (-0.041, 0.019) | N/A | 11668 |  |
| Linear model with no intercept | 0.054 (0.052, 0.056) | Constrained to be 0 | N/A | 11666 | $P = 0.47$ |
| Exponential model | N/A | N/A | 0.00041<br>(0.00039, 0.00043) | 12432 | N/A |

**Figure 4.**
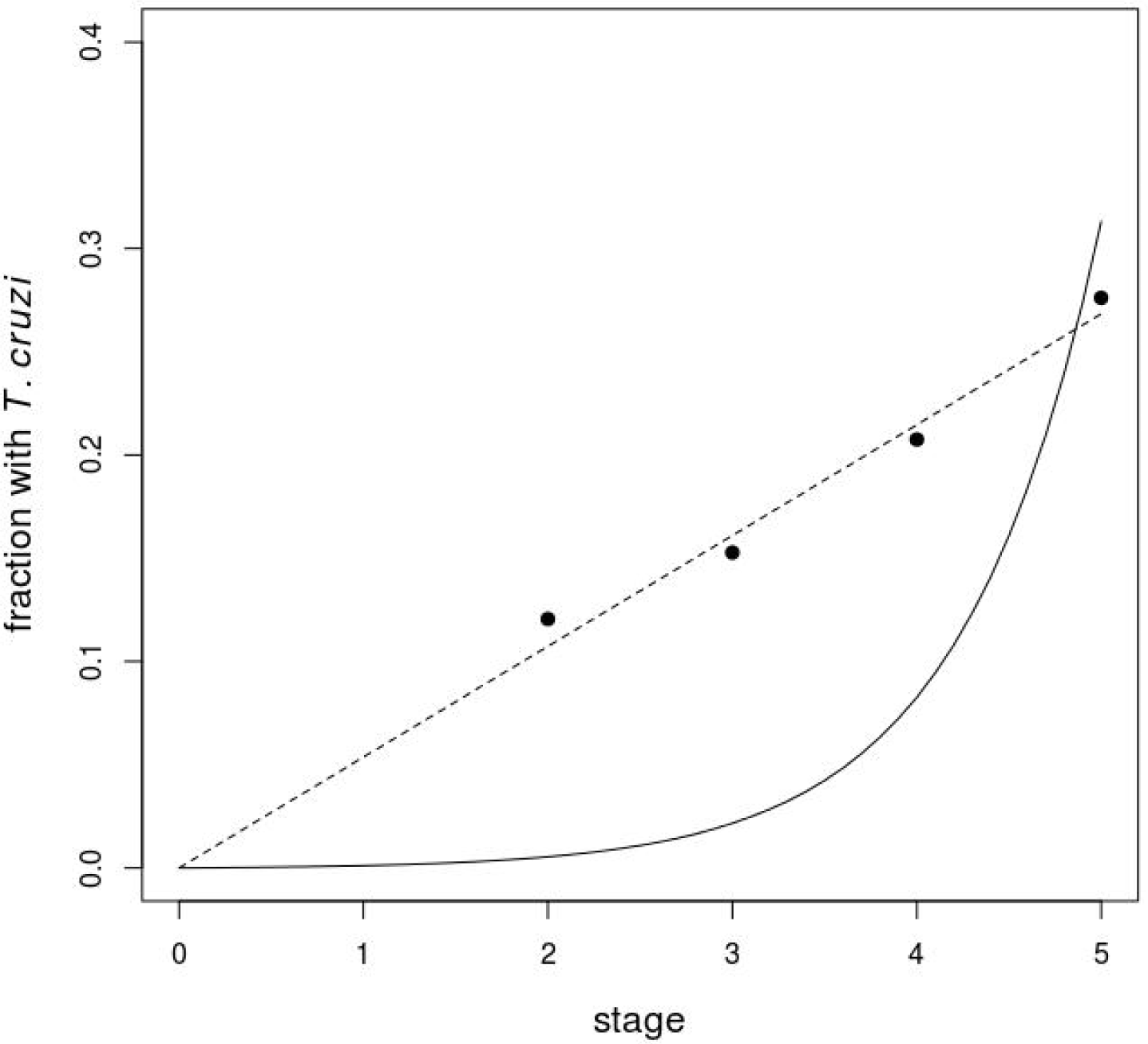
Fraction of triatomines infected with *Trypanosoma cruzi* as a function of stage. Filled circles: mean values observed in 15,252 insects captured at 188 infected sites. Dashed line: simple linear regression model. Solid line: nonlinear regression of infection status on an exponential function of stage.

## 4. Discussion

In a large field survey of *Triatoma infestans* captured in Arequipa, Peru, we demonstrate that *Trypanosoma cruzi* prevalence increases linearly with developmental stage. This result suggests that acquisition of *T. cruzi* depends on the number of feeding opportunities (i.e., bites), and not on the quantity of blood ingested. A possible explanation for this finding is that newly infected mammalian hosts may undergo a rapid logistic increase in the number of circulating parasites. In other words, hosts may go from no parasitemia to a high level of infectiousness very quickly [12,13]. Vectors that feed on such hosts may be almost certain to acquire the parasite, regardless of the size of the blood meal they ingest. Hosts may spend only a brief amount of time at intermediate levels of parasitemia during which vectors acquire the parasite in proportion to the amount of blood ingested. Although this interpretation assumes that instar stage is proportional to the number of blood meals, we note that the exact relationship between these two quantities is uncertain. Some nymphs may take only one blood meal per molt, while others may take multiple blood meals per developmental stage, particularly if their feeding is interrupted before engorgement [14,15].

Our results are in general agreement with those of a prior study that reported increasing *T. cruzi* prevalence with stage in a smaller sample of wild-caught *T. infestans* nymphs and adults [16]. As in the previous study, a vector control program was ongoing in Arequipa during the time period when we collected insects. These control efforts reduced transmission of *T. cruzi* to humans [8] and likely decreased the fraction of infected vectors. We also confirm the results of a previous small study (n <200) of laboratory-reared fifth instar triatomines used for xenodiagnosis, in which there was no correlation between blood meal size and the probability of *T. cruzi* infection [17].

Several reports have cast doubt on the importance of coprophagy as a route of *T. cruzi* acquisition [12,18,19]. The present study provides complementary epidemiologic evidence, from *T. infestans* captured in natural habitats, against the coprophagy route of transmission. However, since we were unable to examine first instars, more work is needed to quantify the precise role, if any, of coprophagy. It should also be noted that triatomines’ need for bacterial symbionts and coprophagy have not been established with certainty. Wild-caught triatomines harbor diverse gut bacteria [6], which suggests they can acquire symbionts from routes other than coprophagy, and many triatomine species (including *T. protracta, T. rubida*, and *Rhodnius prolixus*) develop normally when raised in sterile environments [20].

Limitations of this study include our inability to measure the infection prevalence among first instars; our averaging of the infection prevalence across all sites, which may have oversimplified a more complex pattern of parasite transmission such as may occur in a metapopulation; and uncertainty in our assumptions regarding the number and size of blood meals consumed by triatomines of different stages. Another limitation is that migration of infected triatomines could have biased our prevalence estimates by causing us to include colonies without well-established stable patterns of infection versus stage.

The linear increase in *T. cruzi* prevalence with stage could have implications for Chagas disease control strategies. Insecticides and biological control methods that preferentially kill older insects have relatively less effect on an insect’s reproductive fitness and, in theory, are less likely to be rendered obsolete by evolution of resistance mechanisms [21]. Late-acting agents have been suggested as a means to combat infections including malaria [22] and Chagas disease [23] Had we observed an exponential increase in *T. cruzi* prevalence with stage, with few infected early-stage nymphs, we could have made a strong case for such late-acting methods. Instead, we found non-trivial infection prevalence in second and third instars. Although our results suggest that disproportionate killing of older triatomines may not be useful, these findings should be taken with a grain of salt. The probability that *T. cruzi* will be transmitted to humans depends upon several factors that we did not measure. Behavioral and physiological factors may make younger triatomines less likely to transmit the parasite. Younger instars may be less likely than adult insects to defecate on hosts [24]; may defecate longer after taking a blood meal [25]; and may be less likely to survive the parasite’s incubation period, since survival of infected triatomines appears to be affected by the ratio of ingested blood to body weight [26]. Further studies will be needed to determine the relative risk of feces-mediated transmission posed by younger and older triatomines, which in turn would inform the decision of whether to pursue late-acting control strategies.

## Acknowledgments

We would like to thank the following members of the Zoonotic Disease Laboratory in Arequipa who assisted with this work: Jenny Ancca, Carlos Condori, and Jesus Pinto Caballero. We gratefully acknowledge the invaluable contributions of the Ministerio de Salud del Perú (MINSA), the Dirección General de Salud de las Personas (DGSP), the Estrategia Sanitaria Nacional de Prevención y Control de Enfermedades Metaxenicas y Otras Transmitidas por Vectores (ESNPCEMOTVS), the Dirección General de Salud Ambiental (DIGESA), the Gobierno Regional de Arequipa, the Gerencia Regional de Salud de Arequipa (GRSA), the Pan American Health Organization (PAHO/OPS) and the Canadian International Development Agency (CIDA).

## Appendix A

Rabinovich [4] found that *Triatoma infestans* blood meal sizes increased from 3.6 mg in first instars to 360.5 mg in fifth instars (Table A.1). We used these data to estimate the cumulative amount of blood ingested by captured wild *T. infestans*. We assumed that each insect took one blood meal per stage and that captured insects had recently fed. In other words, prior to its capture, each insect had already consumed the blood meal corresponding to its current developmental stage. Under these assumptions, cumulative blood ingested is equal to the sum of the sizes of the blood meals consumed during the current and all prior stages (Table A.1, column 3)

**Table A.1.** Estimates of blood meal sizes and cumulative blood ingested by *Triatoma infestans* nymphs. Blood meal sizes, in column two, are taken from Rabinovich [4]. Cumulative blood calculations (column 3) assume insects took one blood meal per stage and were recently fed when captured.

| Stage | Size of blood meal<br>(mg) | Cumulative blood<br>ingested during lifetime<br>(mg) |
| --- | --- | --- |
| First instar | 3.6 | 3.6 |
| Second instar | 13.3 | 16.9 |
| Third instar | 45.4 | 62.3 |
| Fourth instar | 174.4 | 236.7 |
| Fifth instar | 360.5 | 597.2 |

We used nonlinear regression to model the cumulative blood ingested as an exponential function of stage. The model was a near-perfect fit to the data (Figure A.1). The best-fit model was

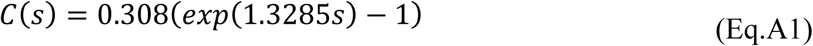

where C is the cumulative blood ingested during the insect’s lifetime and s is the developmental stage.

If acquisition of *T. cruzi* is proportional to cumulative blood ingested, then by definition

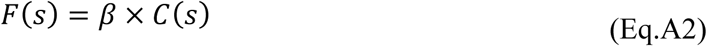

where F is the fraction of insects with *T. cruzi*; β is an unknown proportionality coefficient; and C(s) is cumulative blood ingested as defined above. Substituting Equation (A1) into Equation. (A2) gives

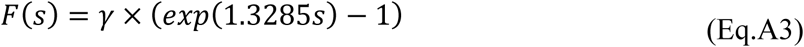

where γ is the proportionality coefficient obtained by combining the other two coefficients (i.e., γ=0.308×β).

**Figure A.1.**
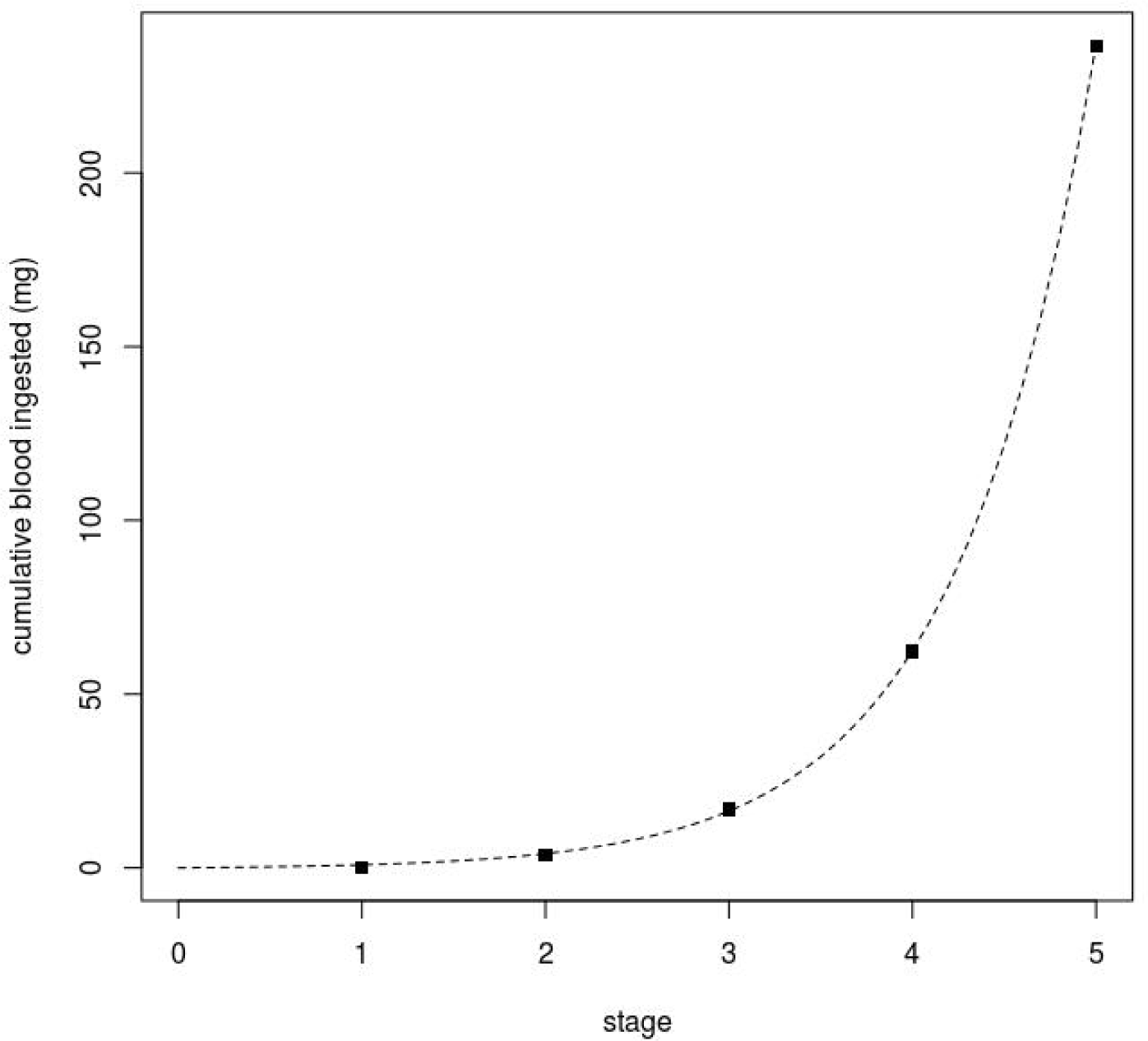
Cumulative blood ingested by *Triatoma infestans* as a function of stage. Black squares are estimates based on blood meal sizes from Rabinovich [4]. Dashed line shows an exponential fit obtained with nonlinear regression.

This result suggests that if *T. cruzi* prevalence depends on cumulative blood ingested, then prevalence should increase exponentially with stage, with a growth constant of roughly 1.3285. In the main text of the manuscript we test this hypothesis by performing a nonlinear regression of *T. cruzi* infection status on the exponential growth function given in equation (Eq.A3).

